# Nanotopography modulates intracellular excitable systems through cytoskeleton actuation

**DOI:** 10.1101/2022.06.09.495528

**Authors:** Qixin Yang, Yuchuan Miao, Parijat Banerjee, Matt J. Hourwitz, Minxi Hu, Quan Qing, Pablo A. Iglesias, Peter N. Devreotes, John T. Fourkas, Wolfgang Losert

**Affiliations:** Department of Physics, University of Maryland, College Park, MD 20742; Institute of Physical Science and Technology, University of Maryland, College Park, MD 20742; Department of Cell Biology, Johns Hopkins University, Baltimore, MD 21205; Department of Physics & Astronomy, Johns Hopkins University, Baltimore, MD 21218; Department of Chemistry & Biochemistry, University of Maryland, College Park, MD 20742; Center for Biological Physics Center, Arizona State University, Tempe, AZ 85281; Department of Physics, Arizona State University, Tempe, AZ 85287; Biodesign Institute, Arizona State University, Tempe, AZ 85287; Department of Electrical & Computer Engineering, Johns Hopkins University, Baltimore, MD 21218

## Abstract

Cellular sensing of most environmental cues involves receptors that affect a signal-transduction excitable network (STEN), which is coupled to a cytoskeletal excitable network (CEN). We show that the mechanism of sensing of nanoridges is fundamentally different. CEN activity occurs preferentially on nanoridges, whereas STEN activity is constrained between nanoridges. In the absence of STEN, waves disappear, but long-lasting F-actin puncta persist along the ridges. When CEN is suppressed, wave propagation is no longer constrained by nanoridges. A computational model reproduces these experimental observations. Our findings indicate that nanotopography is sensed directly by CEN, whereas STEN is only indirectly affected due to a CEN-STEN feedback loop. These results explain why texture sensing is robust, and acts cooperatively with multiple other guidance cues in complex microenvironments.

**One-Sentence Summary:** Cells sense nanotopography directly through their cytoskeletal dynamics, in a reversal of traditional signaling pathway hierarchies.

---

Cells sense and adapt to the physical properties of their environment, such as rigidity (*1, 2*), adhesivity (*3*), and nanotopography (*4, 5*). Sensing of the physical environment is required for many functions of normal cells, including immune response (*6*), proliferation (*7*), and stem-cell differentiation (*8, 9*). Previous research has identified the critical roles of integrins in sensing local adhesivity and rigidity (*3, 10–13*). However, the mechanisms of the sensing of nanotopography are poorly understood.

Nanotopography is an important feature of the microenvironment, and can guide cell morphology and migration by aligning cellular protrusions (*5, 14, 15*). Nanotopography also alters and guides waves of actin polymerization in a wide range of cells, including *Dictyostelium discoideum* (*5*), neutrophils (*16*), and epithelial cells (*16*). In contrast to the two-dimensional waves observed in cells on flat substrates, cells on nanoridges exhibit one-dimensional waves that are guided along the ridges, in a process known as esotaxis. Recent studies have shown that changing the actin-wave patterns leads to transitions in protrusion profiles and migrational modes (*17, 18*).

Here we examine the role of actin waves in sensing nanotopography. Waves of actin polymerization involve a cytoskeletal excitable network (CEN) that is coupled to a signal-transduction excitable network (STEN). STEN serves as the primary sensing module of external chemical cues, and regulates downstream CEN activity (*19, 20*). The features of protrusions are coordinated by traveling waves of this coordinated STEN-CEN system on the plasma membranes and in the proximal cortical regions (*18, 21*). CEN itself can only display local oscillatory behavior, but provides delayed negative feedback to STEN (*18, 22*). Given that the CEN and STEN signaling systems include membrane-bound molecules, and that the sensing of nanotopography must involve the membrane, nanotopographic cues could alter STEN, CEN, or both. Indeed, sensing of electric fields, another physical guidance cue that affects the cell surface, involves both STEN and CEN (*17, 23*). We follow both STEN and CEN markers in electrofused *Dictyostelium discoideum* on nanoridges. Electrofusion merges tens of *D. discoideum* cells into one cell with a correspondingly larger volume and surface contact area, yet does not change the characteristic scales of actin waves (*24*). Thus, electrofused cells contain a large number of actin waves that are not impacted by the edge of the surface contact area. We apply a variety of acute perturbations to probe the roles of STEN and CEN in sensing of this nanotopography.

## Results

### Patterns of signal transduction and actin polymerization are coordinated on nanotopography

We employed PHcrac-GFP and limE-RFP as fluorescent biosensors of PIP3 and F-actin, which represent STEN and CEN activities, respectively (*18, 22*) in electrofused *Dictyostelium discoideum* cells (Fig. 1A). We found that the activities of F-actin and PIP3 were correlated on both flat surfaces and nanoridges, but with different spatial organizations. On flat surfaces, 2- to 3-μm-wide bands of PIP3 traveled radially on the basal membrane with a speed of 6 to 10 μm /min. F-actin appeared throughout the bands, although higher levels decorated the leading and trailing edges of the PIP3 waves (Fig. 1A, Movie S1). In cells plated on aligned nanoridges (width ~150 nm, height ~100 nm, spacing 5 μm, Fig. 1B, Movie S2), the ridges triggered a pronounced rearrangement of the F-actin and PIP3 wave patterns (Fig. 1C). Strong F-actin fluorescence signals were observed on ridges, but PIP3 fluorescence signals were not enhanced compared to those on a flat substrate (Fig. 1C). F-actin and PIP3 formed traveling waves that filled the valleys between the ridges. Even though the ridge height (~100 nm) was smaller than the height of a typical actin wave (~ 0.8 μm), the nanostructures effectively guided the waves. In Fig. 1C, a wave was generated in the valley between two ridges (0 s). A bright F-actin streak formed when the wave grew and reached the ridge (90 s in fig. 1C). The wave did not propagate across the ridge, but propagation parallel to the ridge continued (90 s to 180 s in Fig. 1C).

**Figure 1.**
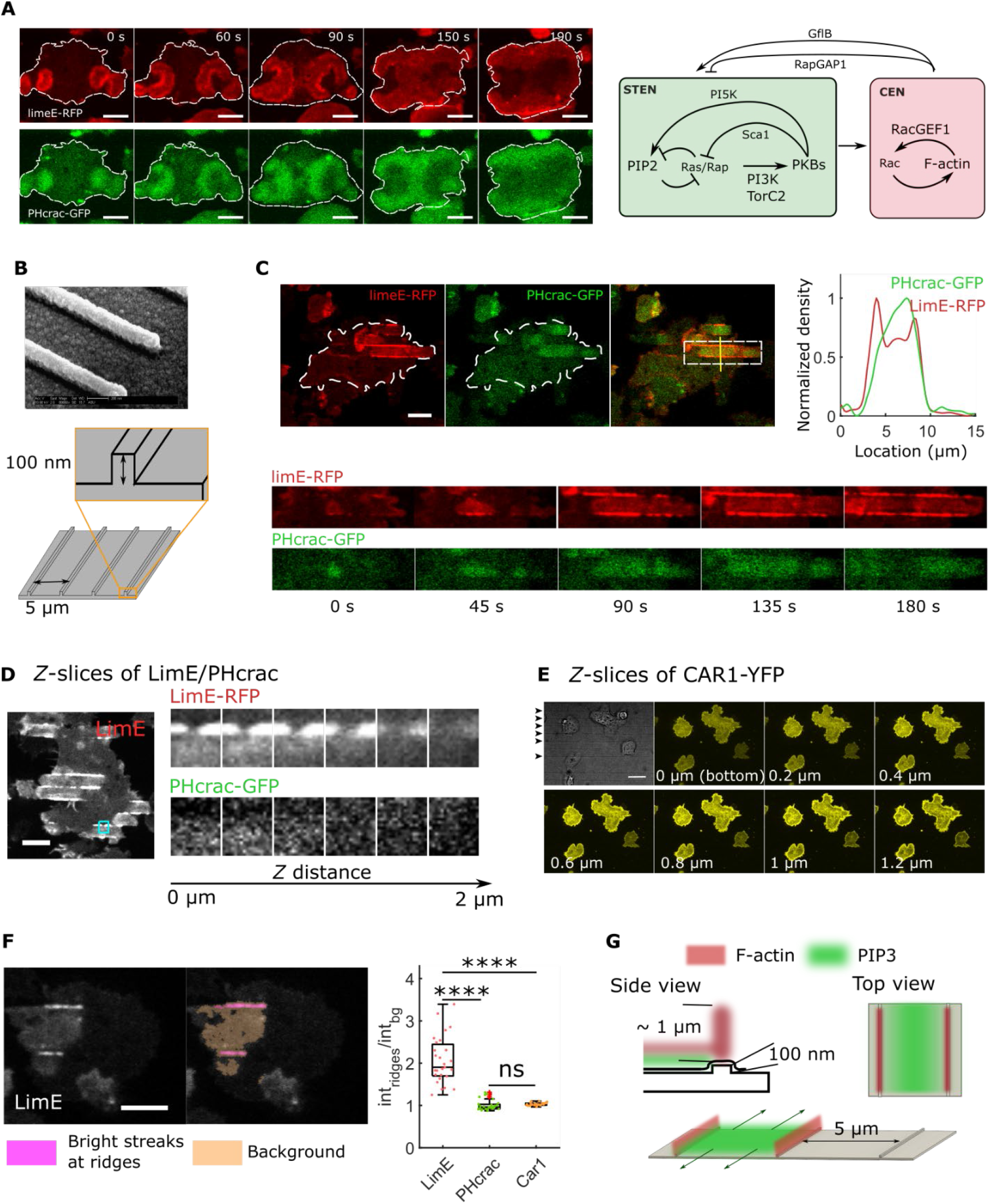
Coordinated activities of STEN and CEN components on flat surfaces and nanotopographies. A. Time-lapse images of limE-RFP (top row) and PHcrac-GFP (bottom row) from an electrofused giant cell on a flat resin surface. LimE-PH waves propagate radially in band-shape, whereas limE signals rim the PH signals. The right schematic shows the current understanding of the STEN and CEN network with key components. B. Shape of the nanoridges. The height of each ridge is 100 nm, and the spacing between neighboring ridges is 5 μm. C. Time series of limE and PHcrac images from the giant cell set on nanoridges shown in B. The profile is from the vertical line in the combined image. The bottom rows show the temporal changes of the rectangular region highlighted in the combined image. D. *Z*-slices of limE-RFP and PHcrac-GFP. E. *Z*-slices of CAR-YFP. F. Fluorescence signals with the LimE/PH/CAR1 signals near ridges (highlighted in pink) divided by those in the surrounding regions (highlighted by orange). The left plot shows the ratios of ridge signals to background signals (*N_LimE_* = 22, *N_PH_* = 22, *N*_*CAR*1_ = 24). The pairwise T-test was implemented for statistical significance. The data were reflect at least three different days of experiments. G. Schematic illustrating the spatial coordination of F-actin and PIP3 in STEN-CEN waves on ridged surfaces. All the scale bars in this figure represent 10 μm.

We also imaged other markers and additional z-planes (Fig. 1D). Signals from limE were visible at higher focal planes (1.6 μm)than were signals from PIP3 (0.8 μm). Fluorescent images of the membrane marker cAR1 showed no pronounced membrane deformation around the ridges (Fig. 1E, 1F), indicating that the enhanced LimE signal at ridges represents an increase in F-actin rather than a membrane fold.

To investigate how the dimensions of nanoridges affect the STEN-CEN response, we plated cells on ridges with a height of 1 μm (Fig. 2A, Movie S3). In contrast to the results on the shorter ridges (Fig. 1), both F-actin and PIP3 signals were enhanced on the taller ridges. PIP3 and F-actin were organized as streaks on both sides of each ridge structure (Fig. 2A). F-actin-PIP3 streaks displayed wave-like behavior, and the wave propagation was guided parallel to the ridges (Fig. 2A). According to the profile, PIP3 was corralled by F-actin, the latter of which extended further towards the valley regions.

**Figure 2.**
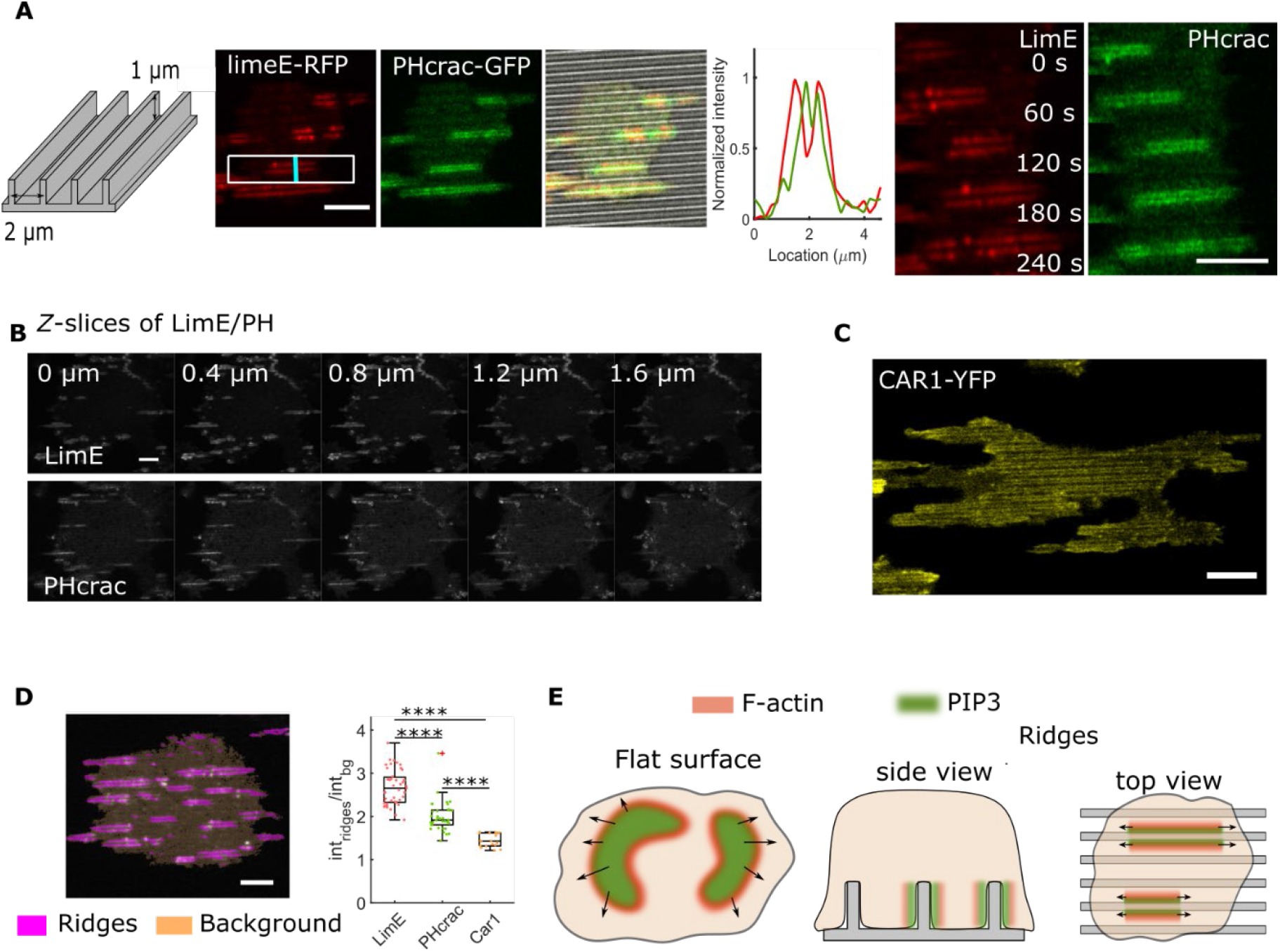
Coordinated activities of STEN and CEN components on 1-μm-high nanoridges. A. LimE (red) and PHcrac (green) images from an electrofused giant cell on 1-μm-high nanoridges with a 2 μm spacing. The profile is from the vertical line in the combined image. The right images show the time lapse of the cropped rectangular box in the limE images. B. *Z*-slices of limE-RFP and PHcrac-GFP. C. Snapshot of CAR-YFP. D. Normalized fluorescence signals. We normalized the LimE/PH/CAR1 signals near ridges (Highlighted by pink) to the background (highlighted by orange). The left plot shows the ratios of the ridge signals to background signals (*N_LimE_* = 40, *N_PH_* = 40, *N*_*CAR*1_ = 33). A pairwise T-test was implemented for statistical significance. The data were collected from at least three different days of experiments. E. Schematic illustrating the spatial coordination of F-actin and PIP3 in STEN-CEN waves on ridged surfaces. All the scale bars in the figure represent 10 μm.

Both F-actin and PIP3 signals were visible up to 1.2 μm above the surface (Fig. 2B). Signals of the receptor cAR1 also showed bright, streak-like structures (Fig. 2C), indicating that the membranes wrap around the taller ridges. The ratio of the fluorescence signals at ridges to the background cytosolic signals from F-actin and PIP3 were stronger than that of cAR1 (Fig. 2D).

Although cAR1 was visible around all ridges covered by a cell, F-actin and PIP3 only appeared around some ridges. These results suggest that although the taller ridges cause membrane deformation, F-actin and PIP3 also respond dynamically to these ridges.

### Altering the STEN states modifies wave patterns

We changed the activation thresholds of STEN by recruiting related proteins to the plasma membranes through chemical-induced dimerization. Inp54p was recruited to the cell membrane to decrease PI(4,5)P2, which led to a higher level of STEN activity. After Inp54p recruitment, the double-streak wave pattern was reduced, and ~40% of the waves became larger in area than the average level under the control conditions. These waves spanned a larger area and crossed multiple ridges (Fig. 3A and 3B, Movie S4). In addition, the average lifetime and propagation speed of the waves both increased (Fig. 3B and 3C, see Method for details).

**Figure 3.**
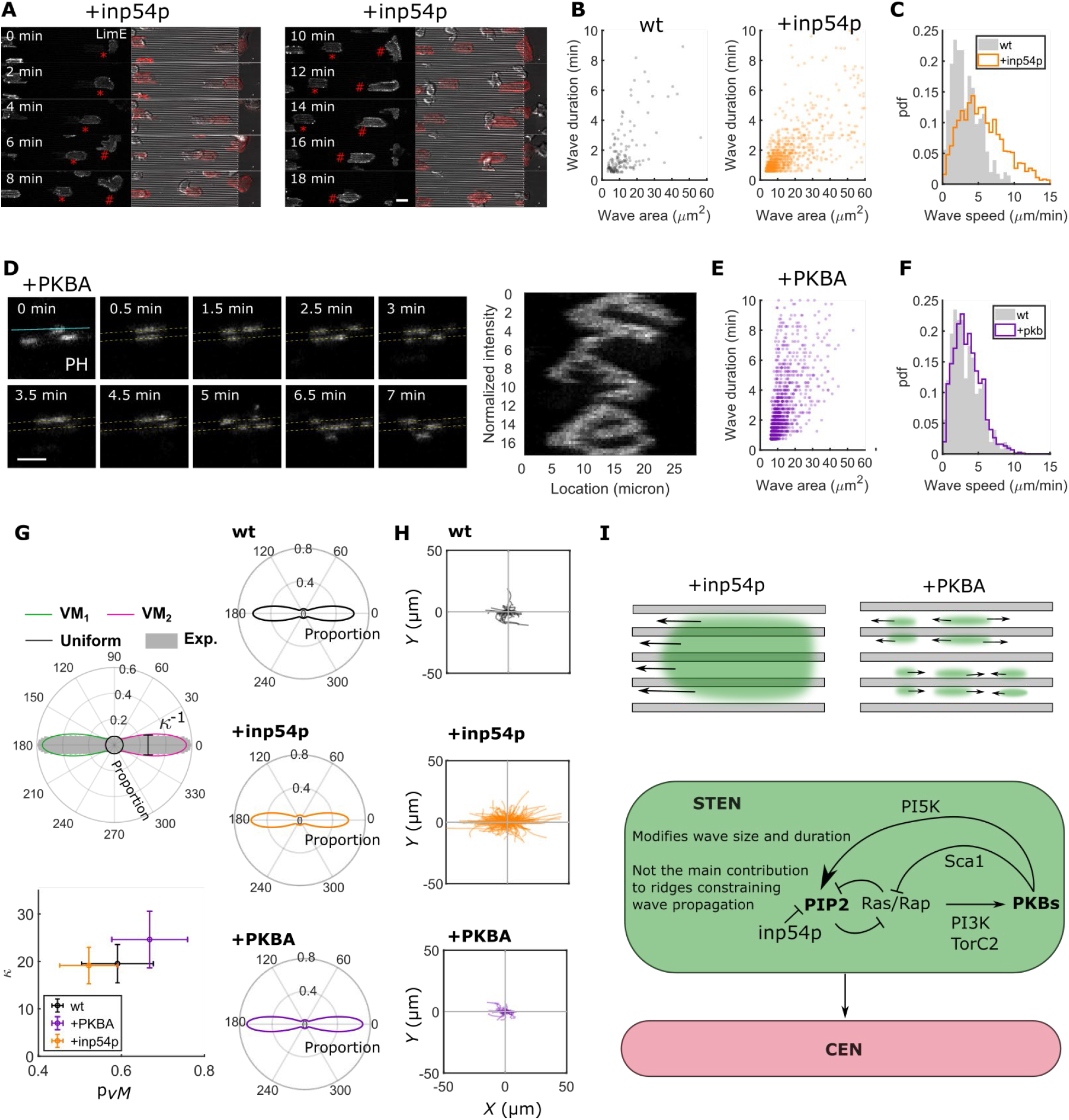
Perturbations on the key nodes of STEN. A. Time-lapse limE images of cells recruited with inp54p. After the inp54p recruitment, waves become larger and span across multiple ridges. B. Scatter plots of wave area vs. wave durations from wild-type and +inp54p experiments. C. Probability distributions of wave propagation speeds for wild-type cells and cells in which inp54p is recruited. D. Time-lapse PHcrac images from cells where PKBA is recruited to the plasma membranes. The kymograph shows the temporal change of PH intensities from the blue line in the image *t* = 0 min. E. Scatter plot of wave area vs. wave duration from cells with PKBA recruitment. F. Probability distribution of wave speed for the +PKBA condition. G. vM model fitting. For each cell, we extract all the reliable optical flow vectors (see Methods for details) and fit the angular to a mixed model with two vM distributions and a uniform circular distribution. In this way we characterize the degree of wave guidance by ridges, using the degree of concentration (*κ*) and the proportion of vMs (*P_νM_* = *P*_*νM*1_ + *P*_*νM*2_). The left bottom plot shows the results of fitted *κ* vs. *PνM*. The right column of polar plots shows the recreated distributions with the average *κ* and average *P_νM_* for each condition. H. Spider plots of tracked cell trajectories for wild-type (top), +inp54p (middle) and +pkba (bottom). I. Schematic showing the changes of wave patterns with different perturbations. The bottom schematic specifies where the chemical perturbations act in STEN and summarizes the role of STEN in sensing nanotopography. All the scale bars in the figure represent 10 μm.

We lowered the activity level of STEN by recruiting PKBA to the plasma membrane (Fig. 3D, Movie S5). Interestingly, we observed that though the wave structures were mostly immobile, they occasionally split into two components that traveled in opposite directions along the ridges (Fig. 3D). The lifetime of the waves increased, but their speed did not change significantly (Fig. 3E and 3F). These observations suggest that wave properties are altered by changing the STEN state, which is consistent with previous studies in which the same perturbations were applied to cells plated on flat surfaces (*18*).

We quantified the guidance of waves using a bimodal von Mises (*νM*) model (see Methods for more details). A previous study found that migration was drastically altered and cells could no longer sense chemoattractants following the Inp54p recruitment (*25*). In contrast, we find that, although the speed and range of the waves increased, the overall guidance of waves (Fig. 3G) and cell migration (Fig. 3H) was not changed significantly by different perturbations to STEN.

### CEN senses nanotopography directly, and nanotopography prolongs the timescale of CEN

In addition to traveling waves of coordinated F-actin and PIP3, stationary F-actin puncta that were independent of PIP3 activity were observed along 1-μm-high ridges (Fig. 4A). Puncta are locations on the cell cortex where actin polymerizes for a brief, characteristic duration without moving laterally. This observation raises the question of whether CEN senses nanotopographic cues directly. To test this hypothesis, we inhibited STEN activity with a combination of PI3K and TORC2 inhibitors, LY294002 and PP242, respectively. After drug treatment, PIP3 activity was absent and actin wave motion ceased. However, bright F-actin puncta were still observed along ridges (Fig. 4B, Movie S6). The localized puncta are consistent with prior findings that cells on flat surfaces retain F-actin puncta following treatment with similar inhibitors (*22*), as we also show in Fig. S1 (Movie S7). F-actin puncta are also observed in the flat regions between ridges (Fig. 4B), with a shorter lifetime (20±16 sec) compared to the puncta generated on ridges (60±45 sec) (Fig. 4D). In these experiments, we used ridges with a 6 μm spacing to distinguish clearly between the F-actin activity at the ridges and between adjacent ridges (Fig. 4B). The same-cell comparison again showed that ridges prolong the lifetime of F-actin puncta. These results suggest that STEN is required for wave propagation (Fig. 4B), but that the nanotopographic signal can be directly detected and transduced by the cytoskeletal network (Fig. 4E).

**Figure 4.**
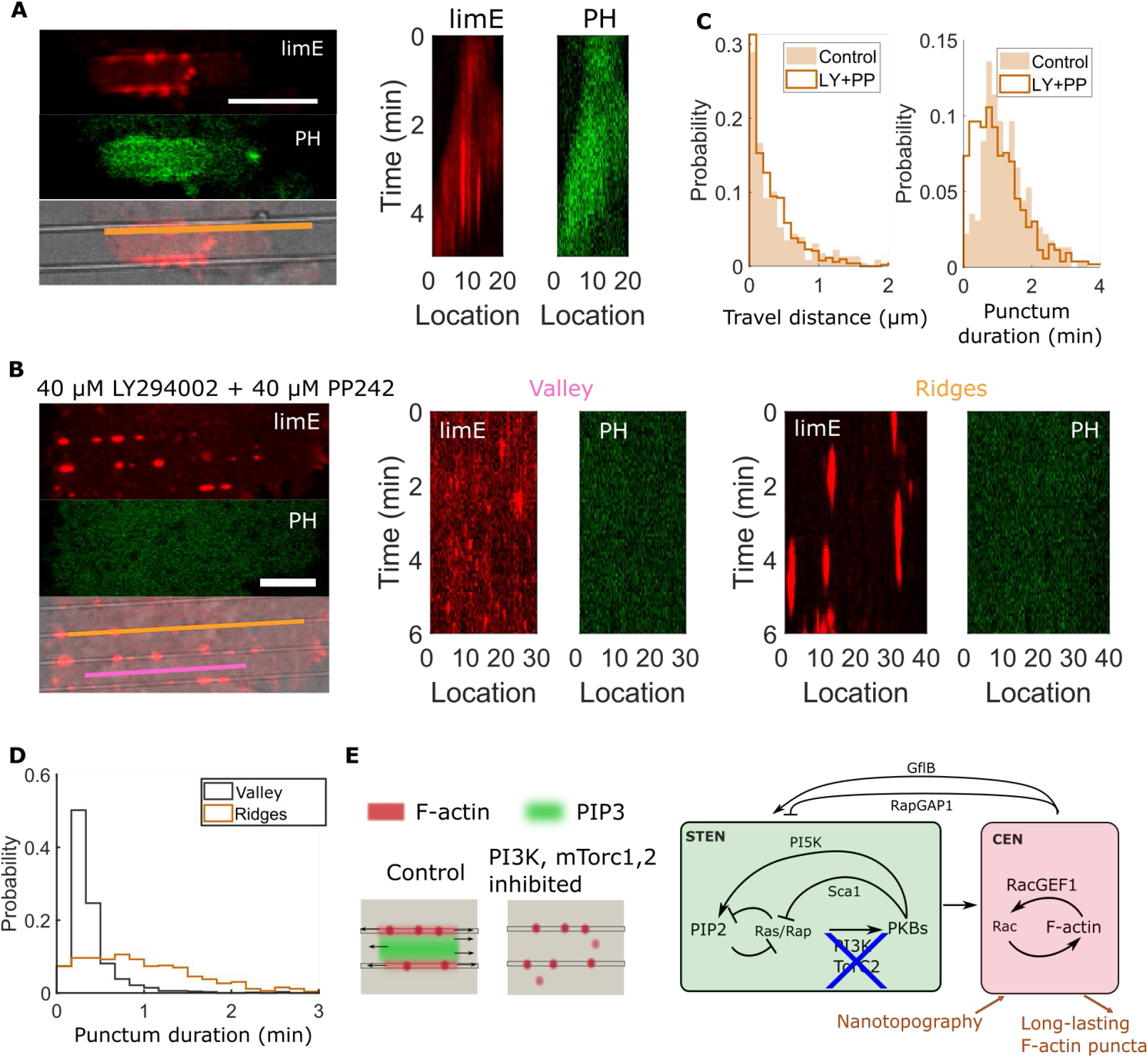
Inhibition of STEN. A. Representative long-lasting F-actin puncta generated near ridges in wild-type cells. The kymographs show the temporal change of signal intensity from the profile in the left image. B. LimE and PH images from a cell treated with LY294002 + PP242 to inhibit PI3K/mTorc1,2 activity. We draw the kymographs from two profiles in the image: the orange line sits on a ridge, and the pink line sits in the valley between two adjacent ridges. According to the kymograph, in both the ridge and valley regions, PIP3 activity is completely inhibited after the drug treatment. F-actin puncta remain, but display distinct durations in the valley and ridge regions. C. Probability distributions of punctum travel distance (*n_LY/PP_* = 530, *n_control_* = 530, two-sample T-test *P* = 10^−4^) and punctum duration (*n*_*LY/PP*_ = 530, *n_control_* = 530, two-sample T-test *P* = 0.0019) in wild-type cells and cells treated with LY+PP242. The data were collected from at least three different days of experiments. D. Comparison of punctum durations for the LY+PP-treated cells on flat surfaces (black) and ridged surfaces (orange). (*n_ridges_* = 705, *n_control_* = 530, two-sample T-test *P* < 10^−10^). E. Schematic illustrating changes of STEN-CEN wave patterns when STEN is inhibited. All the scale bars in the figure represent 10 μm.

### CEN downregulates STEN

Our previous studies have identified negative feedback from CEN to STEN in cells plated on flat surfaces (*18*). We confirmed that this negative feedback also exists in cells plated on nanoridges (Fig. S2, Movie S8). On ridges, RacGEF was recruited to the plasma membrane, PIP3 activity was inhibited (Fig. S2A), and the actin-polymerization level increased (Fig. S2B).

The apparent negative feedback from CEN to STEN (Fig. S2) raises the question of whether nanoridges constrain STEN-wave propagation through enhanced actin polymerization along the ridges. Indeed, when there was an opening in a F-actin streak along a ridge (Fig. 5A), a PIP3 wave often spilled through the opening to reach the neighboring valley. This observation suggests that F-actin serves as a barrier to constrain PIP3 wave propagation (Fig. 5B).

**Fig. 5.**
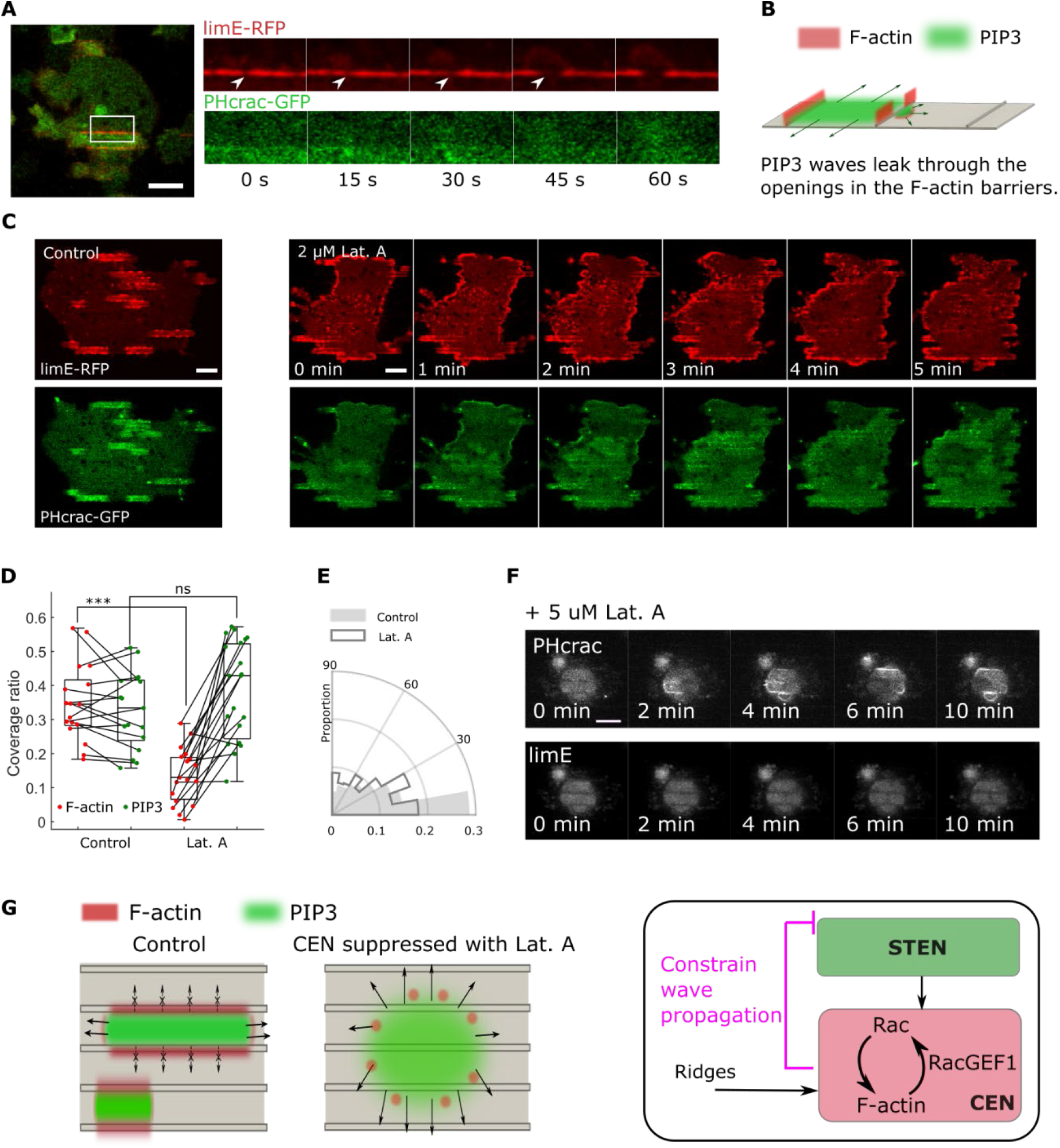
Disrupting F-actin polymerization prevents STEN-CEN wave propagation. A. Time-lapse images showing an F-actin/PIP3 wave that leaks from one valley to another valley on the ridged surface. B. Schematic illustrating how PIP3 waves spread through an opening an F-actin barrier. C. Snapshots of limE-RFP (top) and PHcrac-GFP (bottom) in a giant cell, on ridges, that was treated with 2 μM LatA. D. Comparison of F-actin and PIP3 coverage ratio of cells before/after LatA treatment. Each point represents one giant cell. E. Distribution of wave propagation direction in control and after LatA treatment. F. LimE-RFP (top) and Phcrac-GFP (bottom) images of a giant cell, treated with 5μM LatA, on ridges. G. Schematic illustrating changes in F-actin and PIP3 activity with LatA treatment. All the scale bars represent 10 μm.

To test this hypothesis, we treated cells with a low dose of Latrunculin A (LatA) to suppress, but not fully inhibit, actin polymerization (Fig. 5C, Movie S9). After the addition of 2 μM LatA, actin polymerization decreased (Fig. 5D). The PIP3 levels remained unchanged, but PIP3 waves traveled across ridges, with behavior similar to that on flat surfaces (Fig.5C). We quantified the wave dynamics using optical flow (see Methods). This analysis showed that following LatA treatment, the direction of PIP3 wave propagation was less guided by the ridges (Fig. 5E). We also conducted experiments with 5 μM LatA to inhibit actin polymerization to a greater degree. In this case, PIP3-wave propagation was no longer constrained by the ridges (Fig. 5F). Our observations suggest that the enhanced actin polymerization at ridges serves as a barrier that prohibits PIP3 waves from crossing ridges.

### Results from a computational model of STEN-CEN match experimental findings

We adjusted a computational STEN-CEN-polarity model (*18*) to test our hypothesis regarding the role of nanotopography in the observed wave behavior. This model includes separate, but interconnected, activator/inhibitor excitable networks, representing STEN and CEN, coupled in both a feedforward and feedback manner (Fig. 6A, Methods). The feedback represents polarity, i.e., the tendency of cells to display higher excitation at the location of previous cytoskeletal activity (*18, 26*). In simulations involving flat surfaces, the STEN and CEN waves propagated radially, with the activators leading the respective inhibitors (Fig. 6B). Although STEN activity propagated in a single radial wave, we observed multiple excitations of CEN. The leading wave was solid, but the ensuing waves appeared as puncta. This spatial arrangement is regulated by the different dynamic time scales of the two systems, with STEN being slower and CEN being faster (*18*).

**Fig. 6.**
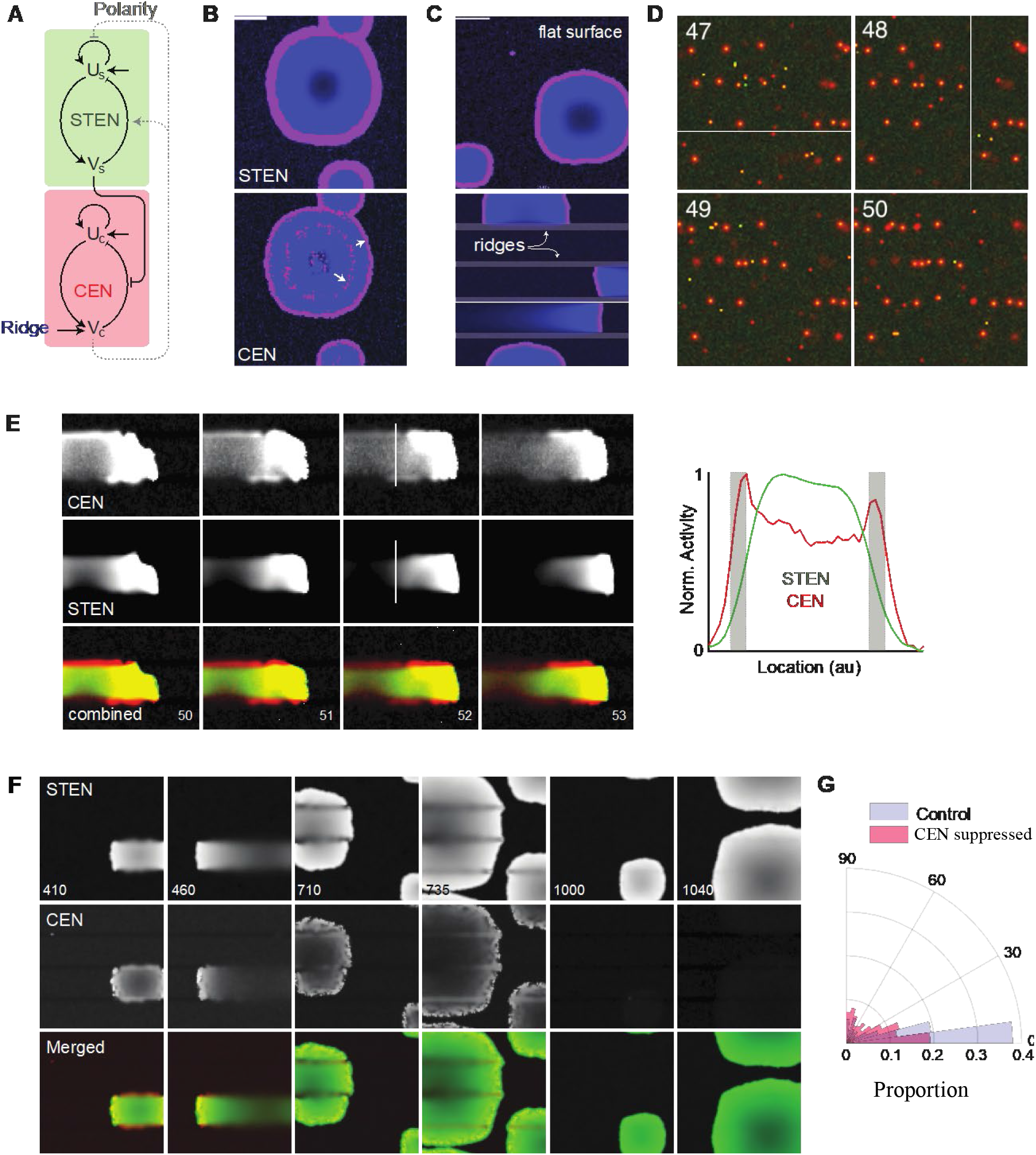
Computational model of STEN-CEN-polarity network recapitulates wave confinement. A. Schematic showing the components of the STEN-CEN-polarity computational model. B. Radial waves in a STEN-CEN-polarity network without ridges. The activators and inhibitors are shown in magenta and blue, respectively. Although a single wave is seen in STEN (top), CEN (bottom) displays multiple concentric excitations, marked by the white arrows. C. Wave patterns of a single EN in the absence (top) and presence (bottom) of simulated ridges. The latter shows the guided propagation along the regions between the ridges. D. Simulations of isolated CEN. The activator (green) and inhibitor (red) co-localize on the ridge as long-lived puncta. E. Simulations of the combined system in which raised CEN activity on the ridges inhibits STEN through the negative feedback of polarity. Although the propagation of both waves is parallel to the ridges, CEN (top and red) waves illuminate the ridges, whereas STEN (middle and green) waves are confined to the valleys. Shown in the right is a line scan of the STEN and CEN intensities along the white line marked in the third column. F. Simulations of latrunculin-treated cells in which CEN activity was steadily lowered in the STEN-CEN-Polarity model with ridges. The waves are confined to the valley initially, but become progressively radial. Eventually, when CEN is inhibited completely, STEN is no longer affected by the ridges. Color scheme as in panel E. G. Distribution of the simulated wave directions in the presence and absence of a fully active CEN network. The polar axis shows the absolute angle of the waves with respect to the alignment of the ridges. The experimental data are presented in Fig. 5E.

To understand how ridges could promote CEN activity and inhibit STEN activity, we investigated a series of increasingly complex network architectures. Although the units of measurement in the simulations were arbitrary, the ratio of the width of the ridge to the width of the valley was set to 1:10, to resemble the pattern of the ridges in experiments. First, we tested how ridges confine wave propagation in a single generic excitable network (EN). In ENs, the activation threshold determines the firing rate and propagation speed (*27*). We conjectured that ridges could direct the waves by raising the local activation threshold. To test this idea, we increased the threshold of a single EN spatially in parallel lines designed to represent ridges. Whereas simulations of the single EN with a uniform threshold showed radial waves (the top image in Fig. 6C), simulations incorporating the elevated threshold showed waves that arose as points in the regions between the ridges, propagated until meeting the ridges, and traveled parallel to the ridges thereafter (the bottom image in Fig. 6C).

We next recreated the effect of ridges on CEN using the simulation model. Experimentally, CEN puncta are longer lasting and brighter on the ridges in cells lacking active STEN (Fig. 4). In simulations of the isolated CEN (Fig. 6D, see Methods for more information), we both lowered the rate of decay of the CEN inhibitor and reduced the CEN threshold on the ridges. The simulations showed behavior similar to that seen experimentally (Fig. 4B, D, and Fig. 6D). Long-lived puncta were located primarily on the ridges, and displayed little propagation. Fewer puncta were observed between the ridges, and those that did appear were of shorter duration than the puncta on ridges.

We then incorporated the effects of ridges on the coupled STEN-CEN-polarity systems (Fig. 6A). We considered two possibilities. First, we assumed that the ridges raise the STEN threshold directly. Although simulations of this scheme directed waves along the valleys (not shown), the model did not reproduce the experimental observations of STEN activity in LatA-treated cells (Fig. 5). Second, we assumed that the higher CEN activity seen at the ridges increases the effect of the CEN-to-STEN feedback, and raises the STEN threshold by simultaneously reducing the positive feedback effect of the STEN activator and increasing the strength of the negative inhibition. Simulations of this scheme successfully recreated both the STEN and CEN wave behaviors on ridges shown in Fig. 1. Whereas the STEN waves propagated longitudinally parallel to the ridges, but were strictly confined to the valleys, CEN waves traveled along the valleys, and wave activity was enhanced on the ridges (Fig. 6E). A cross-section of the corresponding CEN activity shows that in a wave, the activity was high in the valley, peaked on the two ridges, but quickly became negligible on the other sides of the ridges.

Finally, we reproduced the experimental results under F-actin polymerization suppression by LatA (Fig. 5). We simulated a system in which CEN activity steadily decreased from 100% to 1% (Fig. 6F). STEN and CEN waves were confined, and propagated along the valley in the beginning (frames 410 and 460 in Fig. 6F). As the CEN activity decreased, the waves spread to adjacent valleys, and were more radial in appearance (710 and 735). Although CEN was enhanced at the ridges, STEN was absent, as seen in the dark bands on the ridges. Moreover, STEN waves were more diffusive than CEN waves, and CEN waves appeared to be broken and punctate. Finally, as CEN activity disappeared completely (after frame 1000), the STEN waves become radial, which is reminiscent of the simulations on flat surfaces. Optical-flow analysis of STEN waves in the presence or absence of CEN activity showed a greater degree of directed propagation when CEN was active (Fig. 6G), consistent with the analysis of the experimental results (Fig. 5E).

## Discussion

Our work shows that nanoridges modulate STEN-CEN wave behaviors (Fig.1, Fig.2). We applied a variety of acute perturbations to reveal the STEN-CEN components that are key to sensing nanoridges. For most other microenvironmental cues, STEN serves as the entry point and regulator of downstream CEN activities. We showed that STEN is essential for organizing the long-range STEN-CEN wave behavior on flat and ridged surfaces (Fig. 3), but that CEN itself senses nanotopographic cues, with longer-lasting F-actin puncta nucleating at ridges (Fig. 4). We further showed that F-actin serves as a barrier that constrains cross-ridge wave propagation. Suppressing actin polymerization disrupts F-actin barriers, leading to decreased guidance of STEN waves on nanoridges (Fig. 5). Our results suggest that nanoridges modulate STEN wave behaviors through negative feedback from CEN to STEN. This hypothesis is consistent with the results of a well-established computational model of the coupled excitable STEN-CEN system that was adapted to incorporate sensing of nanotopography only in the CEN network (Fig. 6).

### Nanotopography actuates CEN directly

The fact that nanotopography can be sensed directly by CEN distinguishes nanotopography sensing from sensing of most other directional cues, including chemical gradients, the latter of which are perceived via STEN (*19, 20*). Actin polymerization may be affected by the membrane curvature generated by contact with nanotopography through a range of membrane-bound molecules, as outlined in a recent review (*28*). Membrane curvature is well known to modulate the binding of curvature-sensing proteins, such as BAR domain proteins (*29, 30*), which is beyond the scope of the current study, but is worthwhile to investigate in the future. The exact mechanism of enhanced actin polymerization around nanoridges may also depend on cell type, cell state, and the size and shape of the nanoridges.

Although CEN acts as a direct sensor of nanotopography, the coupling of CEN to STEN is essential for using preferential locations of actin polymerization to create guided waves that drive directional motion (*5*), as indicated by our PI3K/mTORC1,2 inhibition results (Fig. 4). This wave-guidance effect of nanotopography is observed in a wide range of cell types on nanoridges (5, 14, 32, 33). When STEN is inhibited, CEN generates oscillating puncta of F-actin, in agreement with previous studies with cells on flat surfaces using the same drug treatment (*18, 22*). The F-actin puncta generated on nanoridged surfaces display a longer lifetime than those generated on flat surfaces, indicating that nanotopographic cues stabilize F-actin (Fig. 4). This experimental observation was successfully recapitulated in our simulations by lowering the CEN inhibitor’s decay rate on the ridges (Fig. 6D).

### Cellular sensing of nanotopography is intermittent due to the dynamic character of the STEN-CEN network

We also plated cells on ridges that were 1 μm high. These ridges induced pronounced membrane curvature (Fig. 2). Both the PIP3 and F-actin biosensor signals are increased on these ridges (Fig. 2A), and the spatial arrangement of PIP3 and F-actin resembles that in phagocytic cups (*31*). Although the membrane is curved around each covered ridge (Fig. 2C), the enhanced F-actin and PIP3 activity only occurs when a wave is formed (Fig. 2A). This phenomenon was observed in previous work, which showed that the response of *Dictyostelium* cells to holes on a perforated film is restricted within the interior regions of STEN-CEN waves (*32*). Our results suggest that membrane curvature does not guarantee a cellular response to nanoridges. Instead, the cell cortex is partitioned dynamically into inactive and active regions (*33*), and actin waves only form in the active area. Nanoridges lower the local activation thresholds. Thus, waves are more likely to be initiated at ridges.

### A mathematical model elucidates the role of feedback from CEN to STEN in the sensing of ridges

In the previously developed STEN-CEN model, STEN senses chemical cues (*26, 34*), or is activated by stochastic noise, before triggering CEN. In chemotaxis, a CEN-to-STEN feedback loop slowly establishes a cell-scale polarity, and gives rise to excitation at locations of prior activity. This cell-scale asymmetry thus provides memory, and increases the response to sensed chemotactic gradients (*26*). Our experimental results indicate that CEN plays a more direct role in sensing nanoridges, and that STEN receives its input from CEN (Fig. 3 and Fig. 5). Our experiments and simulations (Fig. 6) suggest that the coupling from CEN to STEN matches the coupling in chemotaxis, but instead of cell-scale polarity, smaller, ridge-scale, asymmetries appear more quickly, and distinguish ridges from valleys. The primary modification of the model needed to obtain an inhibitory effect on STEN at the sites of the ridges (in agreement with experimental results) was an increase in the strength of feedback from CEN to STEN. The most effective inhibition was achieved by reducing the positive feedback loop of the STEN activator while also increasing STEN degradation rate by the inhibitor.

### STEN waves are guided indirectly by negative feedback from CEN to STEN

On 100-nm-high ridges, F-actin polymerization is enhanced (Fig. 1), whereas PIP3 does not show a direct response (Fig. 1F). However, PIP3 waves do not simply ignore ridges and propagate radially. Instead, wave propagation is substantially constrained by the nanoscale structures (Fig. 1C). Our results in Figs. 5 and S2 suggest that the constraint arises from enhanced F-actin activity, because inhibiting F-actin polymerization leads to cross-ridge propagation of STEN waves. Previous studies hinted that CEN down-regulates STEN, because artificially increasing CEN can eliminate STEN (*18*). However, on flat surfaces, because the timescale of CEN (seconds) usually is much shorter than that of STEN (minutes), this negative feedback has a minor impact on STEN activity. On nanoridges, the timescale of actin polymerization increases to become comparable to that of STEN. The down-regulation effect from CEN to STEN is directly observable as a constraint on STEN-wave propagation by the persistent barriers of polymerized actin that form at the nanoridges. Gaps in the barriers allow STEN waves to spill across ridges, providing strong evidence that the branched cytoskeletal network, and its membrane-bound components, may serve as a diffusion barrier to the STEN signaling lipids and proteins (*35*).

We conclude that the actin cytoskeleton can affect molecules that are generally considered to be upstream of the cytoskeletal machinery. Furthermore, the cytoskeleton is able to influence the localization of nuclear factors YAP (yes-associated protein) and TAZ (transcriptional co-activated with PDZ-binding motif), which leads to shape change of nuclei on nanotopography (*36*). We speculate that microthigmotaxis may involve reversed roles for signaling and the cytoskeleton, and may have distinct downstream effects on the nucleus.

Our work highlights the complementary roles of the two excitable networks that guide cell migration. Whereas STEN is equipped to receive chemical stimuli, our work shows that CEN acts as the primary sensor of nanotopographic guidance cues. Coupling from CEN to STEN is essential in both cases for generating waves that convert preferential locations of polymerized actin into directionally guided waves (esotaxis), and, ultimately, cell migration.

## Acknowledgments

We thank M.Zhao, T. Banerjee, K. Cao, A. Upadhyaya, and W. T. Hill for fruitful discussions. We also thank S. Bhattacharya and D. Biswas for providing assistance in building the simulation model.

## Author contributions

Conceptualization: QY, YM, PB, MJH, MH, QQ, PAI, PND, JTF, WL
Methodology: QY, YM, PB, MJH, MH, QQ, PAI, PND, JTF, WL
Investigation: QY, YM, PB, MJH, MH
Visualization: QY, PB
Formal analysis: QY, PB
Funding acquisition: QQ, PAI, PND, JTF, WL
Project administration: QY, WL
Supervision: QQ, PAI, PND, JTF, WL
Writing – original draft: QY, YM, PB, MJH, MH, QQ, PAI, PND, JTF, WL

## Supplementary Materials

Materials and Methods

Figs. S1-S2

Movies S1-S9

## References and Notes

1. B. L. Doss, M. Pan, M. Gupta, G. Grenci, R. M. Mège, C. T. Lim, M. P. Sheetz, R. Voituriez, B. Ladoux, Cell response to substrate rigidity is regulated by active and passive cytoskeletal stress. Proc. Natl. Acad. Sci. U. S. A. 117, 12817–12825 (2020).

2. L. Trichet, J. Le Digabel, R. J. Hawkins, S. R. K. Vedula, M. Gupta, C. Ribrault, P. Hersen, R. Voituriez, B. Ladoux, Evidence of a large-scale mechanosensing mechanism for cellular adaptation to substrate stiffness. Proc. Natl. Acad. Sci. U. S. A. 109, 6933–6938 (2012).

3. I. D. Campbell, M. J. Humphries, Integrin structure, activation, and interactions. Cold Spring Harb. Perspect. Biol. 3, 1–14 (2011).

4. X. Sun, M. K. Driscoll, C. Guven, S. Das, C. A. Parent, J. T. Fourkas, W. Losert, Asymmetric nanotopography biases cytoskeletal dynamics and promotes unidirectional cell guidance. Proc. Natl. Acad. Sci. U. S. A. 112, 12557–12562 (2015).

5. M. K. Driscoll, X. Sun, C. Guven, J. T. Fourkas, W. Losert, Cellular contact guidance through dynamic sensing of nanotopography. ACS Nano. 8, 3546–3555 (2014).

6. A. Boissonnas, L. Fetler, I. S. Zeelenberg, S. Hugues, S. Amigorena, In vivo imaging of cytotoxic T cell infiltration and elimination of a solid tumor. J. Exp. Med. 204, 345–356 (2007).

7. M. Li, M. J. Mondrinos, X. Chen, M. R. Gandhi, F. K. Ko, P. I. Lelkes, Elastin blends for tissue engineering scaffolds. J. Biomed. Mater. Res. Part A. 79, 963–73 (2006).

8. M. J. Dalby, N. Gadegaard, R. Tare, A. Andar, M. O. Riehle, P. Herzyk, C. D. W. Wilkinson, R. O. C. Oreffo, The control of human mesenchymal cell differentiation using nanoscale symmetry and disorder. Nat. Mater. 6, 997–1003 (2007).

9. J. M. Dang, K. W. Leong, Myogenic induction of aligned mesenchymal stem cell sheets by culture on thermally responsive electrospun nanofibers. Adv. Mater. 19, 2775–2779 (2007).

10. V. Swaminathan, C. M. Waterman, The molecular clutch model for mechanotransduction evolves. Nat. Cell Biol. 18, 459–461 (2016).

11. S. Ghassemi, G. Meacci, S. Liu, A. A. Gondarenko, A. Mathur, P. Roca-Cusachs, M. P. Sheetz, J. Hone, Cells test substrate rigidity by local contractions on submicrometer pillars. Proc. Natl. Acad. Sci. U. S. A. 109, 5328–5333 (2012).

12. S. V. Plotnikov, A. M. Pasapera, B. Sabass, C. M. Waterman, Force fluctuations within focal adhesions mediate ECM-rigidity sensing to guide directed cell migration. Cell. 151, 1513–1527 (2012).

13. J. Z. Kechagia, J. Ivaska, P. Roca-Cusachs, Integrins as biomechanical sensors of the microenvironment. Nat. Rev. Mol. Cell Biol. 20, 457–473 (2019).

14. D. Hoffman-Kim, J. A. Mitchel, R. V. Bellamkonda, Topography, cell response, and nerve regeneration. Annu. Rev. Biomed. Eng. 12, 203–231 (2010).

15. S. Chen, M. J. Hourwitz, L. Campanello, J. T. Fourkas, W. Losert, C. A. Parent, Actin cytoskeleton and focal adhesions regulate the biased migration of breast cancer cells on nanoscale asymmetric sawteeth. ACS Nano. 13, 1454–1468 (2019).

16. R. M. Lee, L. Campanello, M. J. Hourwitz, P. Alvarez, A. Omidvar, J. T. Fourkas, W. Losert, Quantifying topography-guided actin dynamics across scales using optical flow. Mol. Biol. Cell. 31, 1753–1764 (2020).

17. Q. Yang, Y. Miao, L. J. Campanello, M. J. Hourwitz, B. Abubaker-Sharif, A. L. Bull, P. N. Devreotes, J. T. Fourkas, W. Losert, Cortical waves mediate the cellular response to electric fields. eLife 11, e73198 (2022).

18. Y. Miao, S. Bhattacharya, T. Banerjee, B. Abubaker-Sharif, Y. Long, T. Inoue, P. A. Iglesias, P. N. Devreotes, Wave patterns organize cellular protrusions and control cortical dynamics. Mol. Syst. Biol. 15, 1–20 (2019).

19. M. Zhao, B. Song, J. Pu, T. Wada, B. Reid, G. Tai, F. Wang, A. Guo, P. Walczysko, Y. Gu, T. Sasaki, A. Suzuki, J. V. Forrester, H. R. Bourne, P. N. Devreotes, C. D. McCaig, J. M. Penninger, Electrical signals control wound healing through phosphatidylinositol-3-OH kinase-γ and PTEN. Nature. 442, 457–460 (2006).

20. K. F. Swaney, C. H. Huang, P. N. Devreotes, Eukaryotic chemotaxis: A network of signaling pathways controls motility, directional sensing, and polarity. Annu. Rev. Biophys. 39, 265–289 (2010).

21. H. Zhan, S. Bhattacharya, H. Cai, P. A. Iglesias, C. H. Huang, P. N. Devreotes, An Excitable Ras/PI3K/ERK signaling network controls migration and oncogenic transformation in epithelial cells. Dev. Cell. 54, 608–623.e5 (2020).

22. C. H. Huang, M. Tang, C. Shi, P. A. Iglesias, P. N. Devreotes, An excitable signal integrator couples to an idling cytoskeletal oscillator to drive cell migration. Nat. Cell Biol. 15, 1307–1316 (2013).

23. A. L. Bull, L. Campanello, M. J. Hourwitz, Q. Yang, M. Zhao, J. T. Fourkas, W. Losert, Actin dynamics as a multiscale integrator of cellualr guidance cues. Front. Cell Dev. Biol. 10, 2296–634X (2022).

24. G. Gerisch, M. Ecke, R. Neujahr, J. Prassler, A. Stengl, M. Hoffmann, U. S. Schwarz, E. Neumann, Membrane and actin reorganization in electropulse-induced cell fusion. J. Cell Sci. 126, 2069–78 (2013).

25. Y. Miao, S. Bhattacharya, M. Edwards, H. Cai, T. Inoue, P. A. Iglesias, P. N. Devreotes, Altering the threshold of an excitable signal transduction network changes cell migratory modes. Nat. Cell Biol. 19, 329–340 (2017).

26. P. Devreotes, C. Janetopoulos, Eukaryotic chemotaxis: Distinctions between directional sensing and polarization. J. Biol. Chem. 278, 20445–20448 (2003).

27. S. Bhattacharya, P. A. Iglesias, Controlling excitable wave behaviors through the tuning of three parameters. Biol. Cybern. 113, 61–70 (2018).

28. M. M. Kessels, B. Qualmann, Interplay between membrane curvature and the actin cytoskeleton. Curr. Opin. Cell Biol. 68, 10–19 (2021).

29. H. Y. Lou, W. Zhao, X. Li, L. Duan, A. Powers, M. Akamatsu, F. Santoro, A. F. McGuire, Y. Cui, D. G. Drubin, B. Cui, Membrane curvature underlies actin reorganization in response to nanoscale surface topography. Proc. Natl. Acad. Sci. U. S. A. 116, 23143–23151 (2019).

30. S. De Martino, W. Zhang, L. Klausen, H. Y. Lou, X. Li, F. S. Alfonso, S. Cavalli, P. A. Netti, F. Santoro, B. Cui, Dynamic manipulation of cell membrane curvature by light-driven reshaping of azopolymer. Nano Lett. 20, 577–584 (2020).

31. G. Gerisch, M. Ecke, B. Schroth-Diez, S. Gerwig, U. Engel, L. Maddera, M. Clarke, Self-organizing actin waves as planar phagocytic cup structures. Cell Adhes. Migr. 3, 373–382 (2009).

32. M. Jasnin, M. Ecke, W. Baumeister, G. Gerisch, Actin organization in cells responding to a perforated surface, revealed by live imaging and cryo-electron tomography. Structure. 24, 1031–1043 (2016).

33. B. Schroth-Diez, S. Gerwig, M. Ecke, R. Heger, S. Diez, G. Gerisch, Propagating waves separate two states of actin organization in living cells. HFSP J. 3, 412–427 (2009).

34. C. Shi, C. H. Huang, P. N. Devreotes, P. A. Iglesias, Interaction of motility, directional sensing, and polarity modules recreates the behaviors of chemotaxing cells. PLoS Comput. Biol. 9, e1003122 (2013).

35. N. L. Andrews, K. A. Lidke, J. R. Pfeiffer, A. R. Burns, B. S. Wilson, J. M. Oliver, D. S. Lidke, Actin restricts FceRI diffusion and facilitates antigen-induced receptor immobilization. Nat. Cell Biol. 10, 955–963 (2008).

36. Y. Yang, K. Wang, X. Gu, K. W. Leong, Biophysical regulation of cell behavior—cross talk between substrate stiffness and nanotopography. Engineering. 3, 36–54 (2017).

